# Mapping the pathway for protein secretion in the secondary endosymbiotic alga *Nannochloropsis oceanica*

**DOI:** 10.64898/2026.05.25.727594

**Authors:** Tim Michelberger, Anna Santin, Irene Collizzolli, Noor Gammoh, Tomas Morosinotto

**Author notes:** Corresponding authors: Tomas Morosinotto; Anna Santin. **Author Contributions:** Tim Michelberger: Conceptualization, Investigation, Visualization, Writing – original draft. Anna Santin: Investigation, Visualization, Writing – original draft. Irene Collizzolli: Investigation. Noor Gammoh: Writing – review and editing, Supervision. Tomas Morosinotto: Conceptualization, Writing – review and editing, Supervision, Resources. **Competing Interest Statement:** The authors declare that they have no known competing financial interests or personal relationships that could have appeared to influence the work reported in this paper.

## Abstract

Microalgae are key primary producers in marine ecosystems, and their interactions with the surrounding environment rely on the secretion of intracellular metabolites and macromolecules, particularly proteins, supporting essential functions such as nutrient acquisition, environmental sensing and biotic interactions. Most abundant and ecologically relevant seawater algae are secondary endosymbionts, where multiple endosymbiotic events extensively reshaped plastids and intracellular membrane systems, requiring adaptation of protein trafficking mechanisms. This study presents the identification of signal peptides that direct protein secretion in the seawater microalga *Nannochloropsis oceanica*. Their expression in frame with a fluorescent tag enabled to reconstruct the protein secretion pathway in this organism. Proteins channelled for export are first targeted to the periplastidial compartment, an exclusive structure of secondary endosymbiotic algae, that acts as hub for protein trafficking. Subsequently, vesicle-mediated transport directs proteins through the endoplasmic reticulum into the periplasmic space between the cell membrane and the cell wall, from where they are released upon cell division. These findings reveal an evolutionarily remodeled protein secretion pathway, in which host- and endosymbiont-derived trafficking mechanisms merged into an integrated functional system.

**Significance Statement:** The most abundant and ecologically relevant marine algae are secondary endosymbionts whose evolution required extensive re-adaptation of multiple cellular processes. Among them, protein secretion is essential for the interaction with external environment, and required specific re-shaping to the increased cellular complexity associated with endosymbiosis. This work uncovers protein secretory pathway in the secondary endosymbiont seawater alga *Nannochloropsis oceanica* showing that is does not follow a direct route, but proteins are first accumulated in the periplastidial compartment, a unique structure derived from its endosymbiotic history, before being directed for secretion. The final pathway integrated components derived from both the host and endosymbiont, highlighting how evolution was able to merge different biological modules to build an integrated and functional system.

## Introduction

Protein secretion is a crucial mechanism found across all domains of life (1–3), supporting a wide range of physiological functions such as inter- and intra-special interactions (4–6) or sensing of the extracellular environment (7, 8). While prokaryotes lack membrane-bound organelles, meaning that protein secretion involves the direct translocation across their plasma membrane (9), eukaryotes employ complex multi-step secretion systems involving different organelles and pathways for protein processing and sorting before export (8). In photosynthetic microorganisms such as microalgae, secreted proteins plays multiple essential roles: extracellular carbonic anhydrases drive CO_2_ assimilation and regulate pH homeostasis (10, 11), glycoproteins and mucilage components are key elements in the phycosphere and biofilm formation (12), while signalling peptides are responsible for intercellular communication in aquatic ecosystems (13).

The fundamental mechanisms of secretory pathways are supposedly conserved across eukaryotes (3, 14) and mediated by specific signal peptides (SPs), which are genetically encoded at the N-terminus of respective proteins and cleaved during protein export (14). For conventional secretion, nascent proteins with secretory SPs are post- or co-translationally translocated into the endoplasmic reticulum (ER), where the SPs are cleaved off. *Via* vesicular transport, proteins pass through the Golgi apparatus and are eventually released into the extracellular space through exocytosis. Alternatively, non-conventional protein secretion involves protein-filled vesicles directly translocated to the cell membrane, bypassing the Golgi, or proteins simply travelling through the cytoplasm and released by transporters across the cell membrane (15, 16).

Protein secretion routes remain unresolved in secondary endosymbiotic microalgae, whose organelles and endomembrane networks have been extensively reshaped by multiple rounds of endosymbiosis. Such reorganization introduced extra membrane layers, generated unique organelle interfaces, thus requiring modified trafficking routes accounting for additional structural constraints (17, 18). The relevance of this issue is further underscored considering that secondary endosymbiotic algae are the most prominent primary producers in ocean environments and their ecological significance depends on intracellular trafficking for communication with the surrounding environment (19, 20).

Seawater algae of the genus *Nannochloropsis* are secondary endosymbionts belonging to the Stramenopiles lineage, with a chloroplast encased by four membranes (21). The two outermost membranes are the chloroplast endoplasmic reticulum (cER) (22), which is continuous with the host nuclear membrane and endoplasmic reticulum (ER), and the periplastidial membrane (PPM), which has evolved from the former cell membrane of the endosymbiont. The two innermost membranes correspond to the outer and inner envelope membranes (oEM and iEM) of the symbiont chloroplast. The lumen between the PPM and the two innermost membranes of the chloroplast, formerly constituting the cytoplasm of the endosymbiont, forms the periplastidial compartment (PPC) (22, 23), with a putative role in protein quality control, metabolic pathways and plastid division (24, 25). Moreover, *Nannochloropsis* cells are delimited by a membrane and a rigid, multi-layered cell wall, creating a thin periplasmic space in between (26).

In this study, three secretory signal peptides targeting proteins for export in *N. oceanica* were identified and expressed in frame with a fluorescent tag to investigate the protein secretion pathway in this organism. Results show that this starts with protein translocation to the PPC and the subsequent transport across the cER into the periplasmic space, where proteins are released only upon cells division. This pathway shows features inherited from both host and endosymbiont merging into a single functional secretion pathway, as a result of the evolutionary history of the organism.

## Results

### 1 Identification of signal peptides mediating protein secretion in *N. oceanica*

Potential signal peptides (SPs) able to target proteins for secretion in *Nannochloropsis oceanica* were identified by searching for conserved homologs of secreted proteins in available secretomes from related organisms. Putative SPs in *N. oceanica* were predicted for the homologues of the periplasmic metal binding protein TroA (SP1, Figs. 1A, S1A and S2) (27), the glycosyl hydrolase family 16 (SP2, Figs. 1A, S1B and S3) (28) and the cysteine endopeptidase CEP1 (SP3, Figs. 1A, S1C and S4) (29), revealing a high degree of amino acid conservation between SP1 and SP2 (Fig. 1A), and chemical similarity among all three (Fig. 1B). All the three SPs included a positively charged N-terminal residue followed by a hydrophobic patch, which are hallmarks of secretory SPs (30), while only the SP2 contained the frequently found AXA-motif before the predicted cleavage site (Fig. 1A, B).

**Figure 1.**
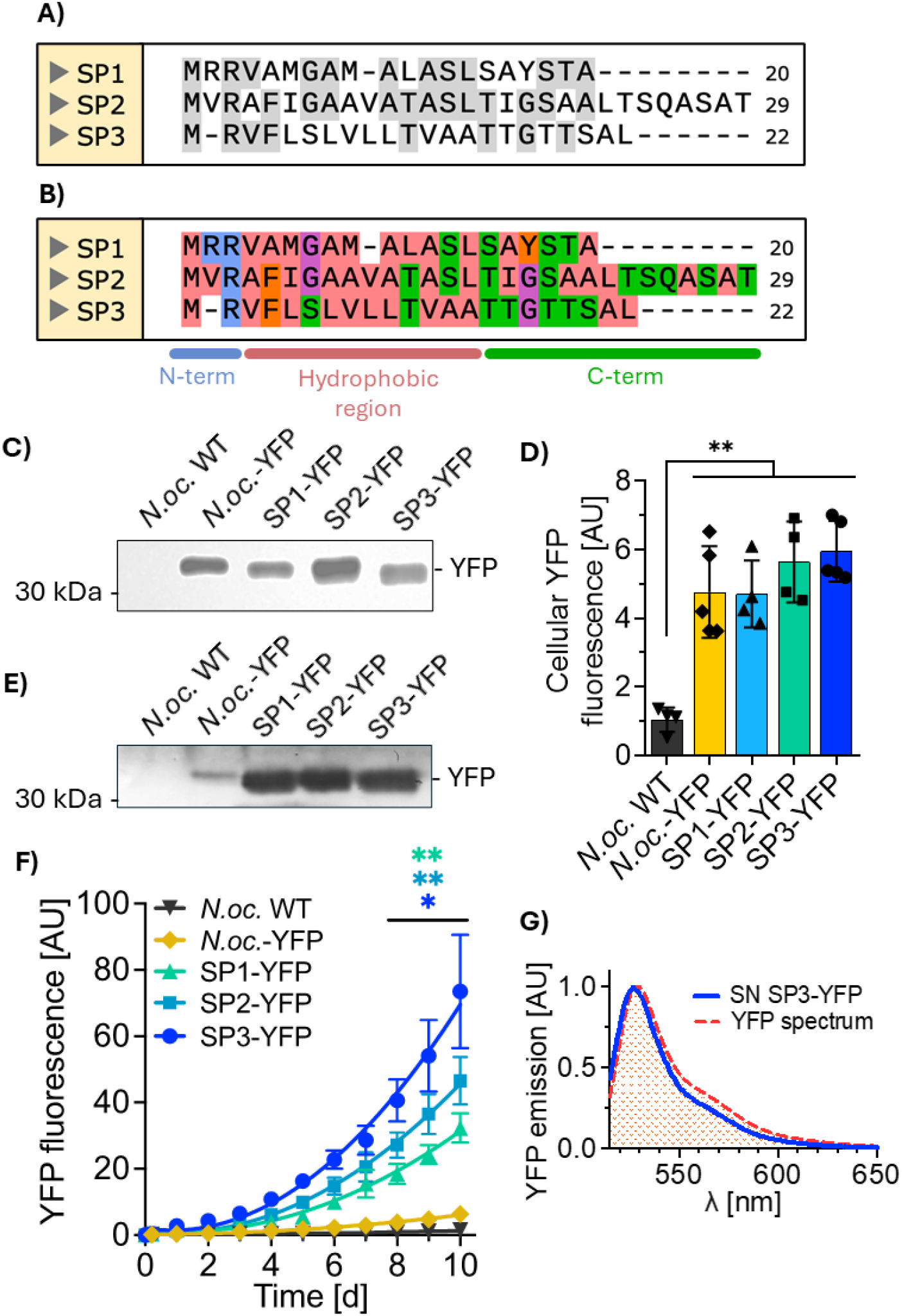
Identification and expression of putative secretory signal peptides in *N. oceanica*. **A)** Sequence similarity between the three putative secretory signal peptides SP1, SP2 and SP3 from *N. oceanica*. Amino acid sequences were aligned with Clustal Ω and processed with SnapGene. Gray shaded amino acid residues mark residues that are conserved in at least another SP. **B)** Aligned sequences from A) were colored after their physico-chemical properties. Blue: positively charged, red/ orange: hydrophobic, purple: small, green: hydrophilic. **C)** Generation of (SP-)YFP-overexpressing *N. oceanica* lines and analysis of (SP-)YFP accumulation. Total cell lysates of *N*.*oc*. WT and randomly selected lines of *N*.*oc*.*-*YFP and SP3-YFP were separated by SDS-PAGE and YFP (27 kDa) or SP3-YFP (31 kDa) were detected by Western blot using α-GFP antibody and run a bit lower than expected. For all lanes, an equivalent of 0.5 µg chlorophyll was loaded. **D)** Cells of *N*.*oc*. WT and randomly selected lines of *N*.*oc*.*-*YFP and the three SP-YFP strains were inoculated in fresh medium at 1.5 x 10^8^ cells ml^-1^. Cell samples of each culture were taken after 3 days, cells were washed twice with fresh medium, and cellular fluorescence was measured with a plate reader. Fluorescence intensity for each measurement was normalized to background fluorescence of the medium. Data is shown as means ± SD from ≥ four independent biological replicates. Significant differences between the lines were assessed with a one-way ANOVA test. **p < 0.01: ns, not significant. **E)** YFP accumulation in the supernatants of the analyzed strains. Samples of all strains were taken on day 7 of the experiment in A), concentrated 50x by centrifugation and analyzed by SDS-PAGE and Western blot. YFP (around 31 kDa) was detected with an antibody against GFP. **F)** Randomly selected lines of *N*.*oc*. WT, *N*.*oc*.*-*YFP, SP1-YFP, SP2-YFP and SP3-YFP strains were inoculated at 1.5 x 10^8^ cells ml^-1^ in fresh medium and YFP fluorescence in the supernatant was measured over 10 days. Fluorescence intensity signal was normalized to background signal of F/2 Plus medium for every individual measurement. Data points represent means ± SEM from at least three independent biological replicates. Curves show fitting to second order polynomial functions. Significant differences of the SP-YFP lines to the *N*.*oc*.*-* YFP line in the last three days were assessed with one-way ANOVA tests for each individual day; *p < 0.05; **p < 0.01. **G)** Representative spectrum of supernatant containing YFP. A sample of the supernatant (SN) of the SP3-YFP line was taken on day 7 from the experiment in A), excited at 515 nm and the fluorescence emission was recorded. The emission spectrum was normalized to the maximum value at the peak at 530 nm. For comparison, the spectrum of YFP was retrieved from fpbase.org and plotted in the same graph.

The three putative SPs identified were genetically fused to a YFP gene (named SP1-YFP, SP2-YFP and SP3-YFP, respectively), and cassettes driving their expression were individually integrated into the genome of *N. oceanica*. The correct expression of (SP-)YFP constructs in cells was verified by Western blot (31), with (SP-)YFP showing the same molecular weight of the control strain expressing YFP with no SP (*N*.*oc*.-YFP, Fig. 1C and S5A). YFP accumulation was confirmed by fluorescence (Fig. 1D).

Accumulation of YFP in the supernatants was detected by Western blot (Fig. 1E and S5B), suggesting that all three putative peptides were able to drive the secretion of YFP from the cells. Consistently, YFP fluorescence signal in the supernatants also was detectable (Fig. 2F), with the SP3-YFP line showing the highest values. A fluorescence emission spectrum of supernatant overlapped with the one of YFP, further validating its presence in the supernatant (Fig. 2G).

**Figure 2.**
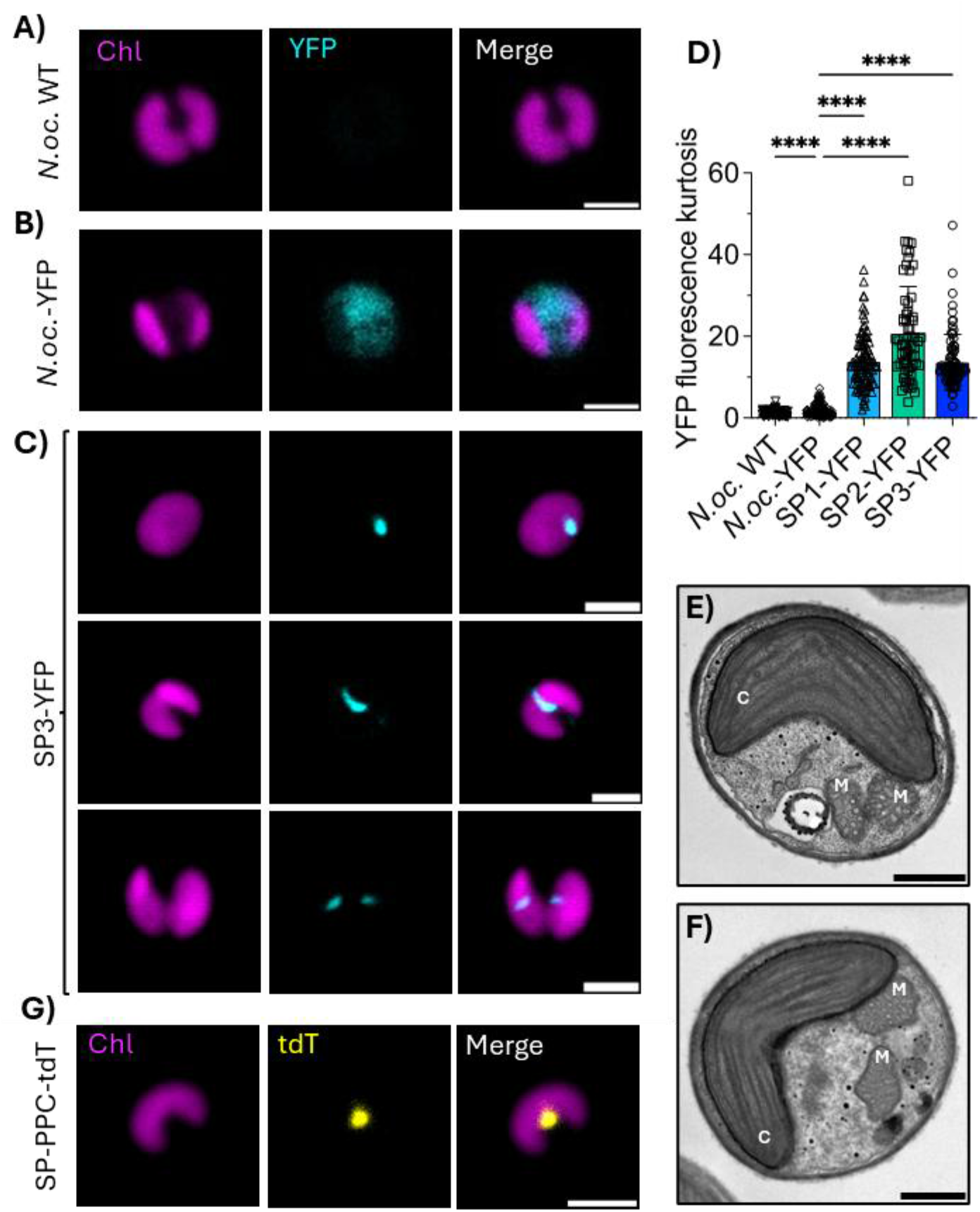
Localization of (SP-)YFP *N. oceanica* lines. **A)** *N*.*oc*. **B)** WT, *N*.*oc*.*-*YFP, and **C)** different morphologies of SP3-YFP cells (during chloroplast division) from cultures grown in shake flasks at continuous illumination with 150 µmol photons m^-2^ s^-1^ were imaged live to analyze sub-cellular localization of YFP. Chlorophyll (Chl) and YFP were imaged by their immanent fluorescence. Scale bars, 2 µm. **D)** Quantification of YFP fluorescence kurtosis from different strains. Cells numbers for both quantifications were n=60 (*N*.*oc*. WT), n=143 (*N*.*oc*-YFP), n=100 (SP1-YFP), n=64 (SP2-YFP) and n=84 (SP3-YFP), and refer to Figures 4 and S10. Data is represented as mean ± SD. Significance was assessed with one-way ANOVA tests; ****p < 0.0001. **E-F)** Electron microscopy of *N*.*oc*.*-*YFP (B) and SP3-YFP (C) cells with dividing chloroplast. Cells from continuous cultures growing at 150 µmol photons m^-2^ s^-1^ were fixed and images of cells with a dividing chloroplast were acquired with electron microscopy. Clearly visible organelles were labelled as: C, chloroplast; M, mitochondrion and N, nucleus. Scale bars, 0.5 µm. **G)** Localization of the PPC in *N. oceanica* in a SP-PPC-tdTomato (SP-PPC-tdT) transgenic line. Chlorophyll (Chl) and tdTomato (tdT) were detected using their immanent fluorescence. Scale bar, 2 µm.

Cell growth of all lines during the secretion experiment, showed no differences between *N*.*oc*. WT and transgenic lines (Fig. S5C), while SDS-PAGE and Coomassie staining on supernatant and cell lysates showed no enrichments of RuBisCo or light-harvesting proteins in the supernatant, highly abundant algal proteins which would be released upon cell death (Fig. S5D). These data confirmed that the high accumulation of YFP in the supernatant was driven by protein export and not by possible toxicity of the (SP-)YFP constructs that could lead to increased cell death and YFP release.

### 2 SP-mediated protein secretion depends on growth conditions

As protein secretion by unicellular organisms is often influenced by environmental parameters (32– 35), growth and YPF fluorescence were monitored for the most effective line, SP3-YFP, exposed to different conditions.

When cultivated at 22°C and continuous light (36, 37) cells showed limited YFP secretion for nine days (Figs. S6A and S7), with a significant YFP export starting from day 10 (Figs. S6A and S8). Intracellular YFP signals were constant throughout the whole experiment (Figs. S6A and S9), suggesting that the delay of YFP export was not limited by the rate of intracellular protein synthesis, but rather determined by a regulation of secretion. At lower temperature, 16°C, cell growth was similar (Figs. S6B and S7), YFP export initiated earlier, reaching similar YFP levels at the end of the experiment. When algal strains were cultivated with a photoperiod of 12:12 hours, thus reducing the total amount of absorbed light by half, this caused slower growth, together with a delayed and reduced YFP export (Figs. S6C, S7 and S8) with no changes were observed in intracellular YFP production (Figs. 3C and S8).

**Figure 3.**
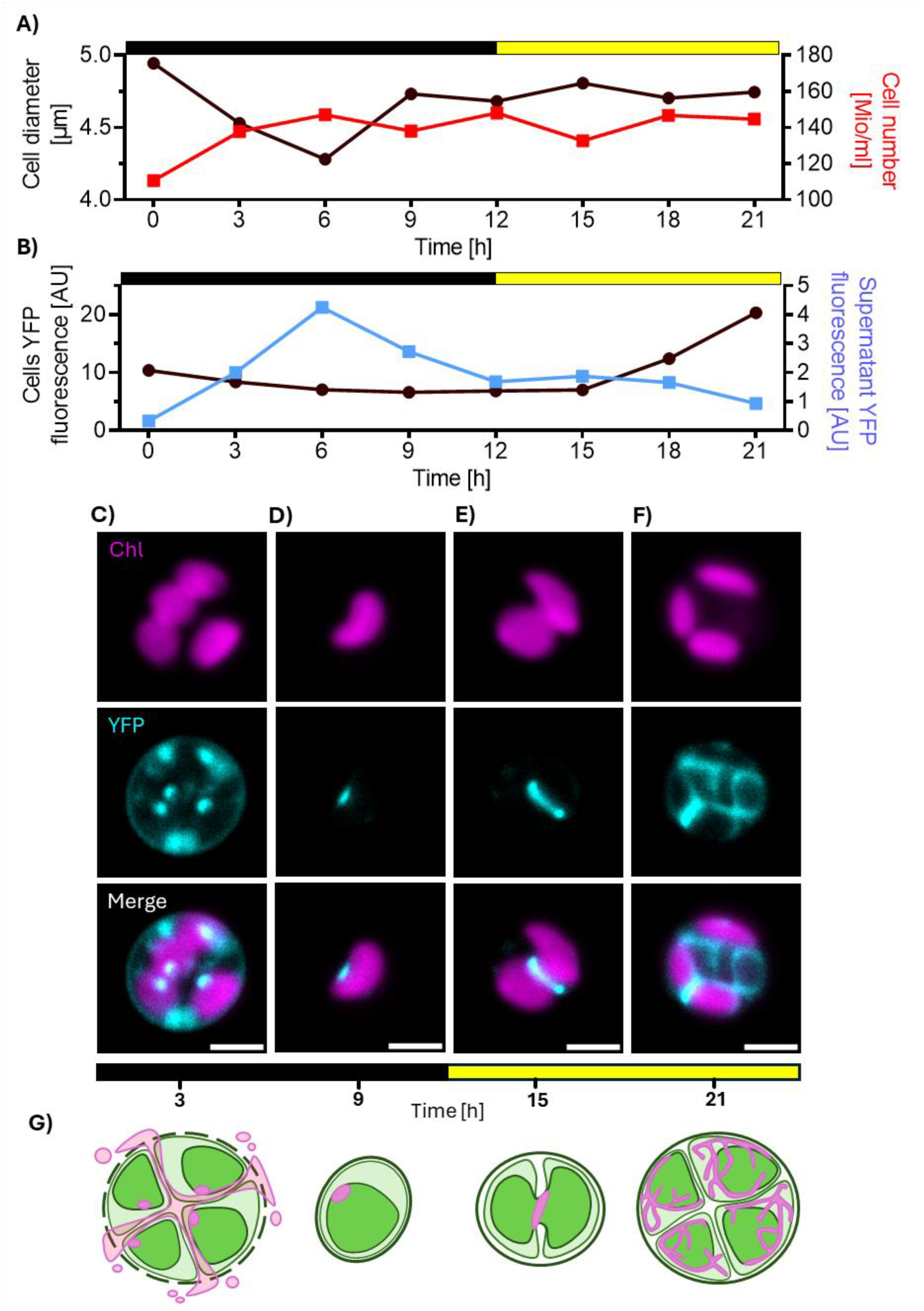
SP3-YFP trafficking over the cell cycle in *N. oceanica* SP3-YFP cells. **A**) Average cell diameter (black) and number of cells (red) in the culture over the cell cycle, determined by a cell counter. **B)** Intracellular YFP fluorescence (black) and YFP fluorescence in the culture supernatant (blue) over the cell cycle. To determine intracellular YFP fluorescence, samples were resuspended in fresh medium for each measurement. Fluorescence signals were normalized to background signal of the fresh medium for each individual measurement. All data points in (A) and (B) represent means ± SD from at least three independent replicates. AU, arbitrary units. **C-F)** SP3-YFP cells observed during the cell cycle. Cells were grown in a 12:12 h light:dark photoperiod with 150 µmol photons m^-2^ s^-1^ during the light phase and an atmosphere containing 1% CO_2_. Cells were resuspended in fresh medium at dusk and samples were imaged 3h (C) and 9h (D) after dusk and 3h and 9h after dawn (corresponding to total 15h (E) and 21h (F)) to follow the subcellular YFP localization over one cell cycle. Chlorophyll (Chl) and YFP were recorded by their inherent fluorescence. Scale bars, 2 µm. means ± SD from three independent replicates. AU, arbitrary units. **G)** Schematic representation of cell cycle and corresponding YFP signal during the process. Pink indicates YFP, dark green for chloroplasts, light green for cytosol.

When cell lines were then cultivated in an atmosphere with 1% CO_2_ to increase carbon availability, and thus photosynthetic activity (38, 39), all strains showed increased growth rates, and reached the stationary phase faster (Figs. S6D and S7). During the exponential growth phase, a massive increase of YFP signal was observed in the supernatant of SP3-YFP, which also plateaued when cells entered the stationary phase (Figs. S6D and S8). The intracellular YFP signal also initially increased during the exponential growth phase, reaching a peak before cells entered the stationary phase (Figs. S6D and S9), and suggesting that the increased secretion might at least partially be attributable to enhanced YFP synthesis. Overall, even if final biomass concentration reached similar values in all tested conditions, YFP levels present in the supernatant were over five times higher when algae were grown with extra CO_2_ supply compared to all other tested conditions, reaching maximum level of 3 mg l^-1^ after 7 days (Fig. S10).

### 3 SPs target YFP to the PPC

To investigate the protein secretion in *Nannochloropsis*, that is an entirely uncharted topic so far, the localization of (SP-)YFP was investigated using confocal laser-scanning microscopy (CLSM). The YFP signal, not detected in *N*.*oc*. WT cells, was broadly distributed in the cytosol in the *N*.*oc*.*-* YFP control cells (Figs. 2A, B and S11A), while it was concentrated in spots or elongated rods in the three (SP-)YFP lines (Figs. 2C and S11A). Such spots or rods that were consistently observed in close proximity to, or in direct apposition with, chloroplasts, as revealed by immediately adjacent chlorophyll autofluorescence (Figs. 2C and S11A). Quantification of the YFP signal distribution confirmed the strong concentration of YFP in the (SP-)YFP strains (Figs. 2D and S11B, C).

(SP-)YFP cells showed different YFP fluorescence patterns during different division stages. In single-chloroplast cells, a single YFP spot was observed, whereas in cells undergoing chloroplast division, the signal extended into an elongated rod-like structure spanning the dividing chloroplasts. Finally, in cells that had completed division, two daughter cells each contained a single chloroplast and a corresponding YFP spot (Fig. 2C). Throughout all stages, the YFP signal was consistently positioned at the chloroplast surface or at chloroplast division sites (Figs. 2C, S11A), not altering chloroplast morphology (Figs. 2E, F and S11D-I).

As secondary endosymbionts, the chloroplast of *Nannochloropsis* cells is wrapped in four membranes of which the two outermost membranes are connected to the ER, creating the Cer (22) while the space between the cER and the inner two membranes of the chloroplast shapes the PPC (22, 23). Fluorescent proteins targeted to the PPC form distinctive spots or slightly elongated structures in both *Nannochloropsis* (40) and the diatom *P. tricornutum* (24), similarly to what was observed in (SP-)YFP cells, suggesting that the proteins targeted for export are trafficked through the PPC. To confirm this hypothesis, a transgenic line expressing a PPC-specific SP (SP-PPC) fused to a fluorescent tdTomato (tdT) protein (40) were generated. The SP-PPC-tdT strain showed the same fluorescent patterns as observed in (SP-)YFP lines, validating the localization of the (SP-)YFP in the PPC (Figs. 2G and S12A). The SP-PPC-tdT strain also showed the same multiple fluorescence patterns, single spots in non-dividing cells and elongated rods in dividing cells, as observed in (SP-)YFP cells, supporting the PPC localization (Figs. 2G and S12A).

To further substantiate this hypothesis, (SP-)YFP cells were examined using markers for other intracellular structures with partially similar morphology in order to exclude alternative localizations. Co-localization analysis with the *N. oceanica* red body, a globular organelle containing ketocarotenoids, was performed by exploiting its intrinsic autofluorescence (Fig. S13A) (41), No spatial overlap was observed between YFP fluorescence and the autofluorescent red body, indicating that YFP does not localize to this structure (Fig. S13A). Nuclei and lipid droplets were visualized using specific fluorescent dyes. Efficient dye penetration occurred only at certain cell cycle stages, likely due to transient changes in membrane permeability, and analyses were therefore limited to adequately stained cells (Figs. S13B, C). Unlike the PPC-associated YFP structures, nuclei were positioned more centrally within the cytosolic space and did not display the tight chloroplast apposition characteristic of the YFP rods (Fig. S13B). Similarly, lipid droplets appeared as multiple cytosolic structures per cell rather than as a single structure associated with each chloroplast, and they showed no consistent spatial correlation with chloroplasts comparable to that observed for the YFP signal (Fig. S13C). Collectively, these observations further support the conclusion that YFP does not localize to the red body, nucleus, or lipid droplets, reinforcing its assignment to the PPC (Fig. 2C).

### 4 Secreted YFP is trafficked from the PPC to the cell envelope *via* the ER

As the YFP localization was shown to be affected by the cells division cycle, a 12:12 hours light:dark photoperiod was applied to synchronize the cells cycle, while also supplying additional CO_2_ to increase protein accumulation and better follow the export pathway. Cells progressively increase in size during the light phase as they undergo successive rounds of internal division, first producing two and subsequently four daughter protoplasts, each containing a single newly formed chloroplast (41, 42). This growth phase is characterized by increased photosynthetic activity, which provides the energetic and metabolic resources required for multiple fission events. However, the resulting four daughter cells remain encapsulated within a single parental cell wall, referred to as the enclosing autosporangial wall, till complete cytokinesis and assembly of their own individual cell walls. Approximately 2-4 hours after dusk, once the individual cell walls of the newly formed daughter cells are fully matured, the autosporangial wall ruptures, leading to the release of four independent, fully differentiated daughter cells (41, 42). This pattern of cell division and release was clearly observed in population dynamics of the synchronized cells. Cell number increased predominantly within the first 6 hours following dusk (Fig. 3A), indicating a synchronized release of daughter cells. This increase in cell concentration was accompanied by a concomitant decrease in average cell diameter (Fig. 3A), consistent with the transition from larger autosporangia to smaller individual cells.

Interestingly, cell multiplication, *ergo* the rupture of the autosporangial wall, also corresponded with an increasing YFP fluorescence in the cell-free supernatant (Fig. 3B) and the decreasing YFP fluorescence in the cells themselves (Fig. 3B). This inverse correlation strongly indicates that YFP, previously accumulated within the cells, was released into the extracellular medium upon autospore rupture and daughter cell release.

When cells were imaged 3 hours after dusk, intact autosporangia were readily identifiable by the presence of four distinct chloroplasts, each corresponding to a newly formed daughter cell enclosed within the autosporangial wall (Fig. 3C). At this stage, YFP presented a fluorescence pattern that exhibited a characteristic punctate pattern on each chloroplast, consistent with SP3-YFP localization to the PPC, as previously observed. Notably, in addition to this expected chloroplast-associated signal, YFP fluorescence was also detected within internal compartments of the autosporangium and was markedly enriched in the cell membrane (Fig. 3C). The YFP localized here is expected to be released at next stage of the cell cycle, namely the rupture of the autosporangium and the release of the daughter cells (Fig. 3A, B). After autospore release, cells did not generate more YFP, likely because of lack of photosynthetic activity in the dark, and the YFP signal remained localized in the PPC (Fig. 3D).

Upon the transition from darkness to light, the cellular YFP progressively increased, suggesting enhanced protein synthesis following the onset of photosynthetic activity (Fig. 3B). Concurrently, chloroplast division was induced during the light phase, as evidenced by the appearance of dividing chloroplasts after 3 hours of illumination (Fig. 3E). At this stage, YFP fluorescence again showed a characteristic sickle-shaped configuration at the chloroplast division site, closely resembling the localization pattern previously observed (Figs. 2C, S12A and 3E).

In these cells, when the light period progressed, YFP fluorescence gradually extended throughout the cell, forming an interconnected network that spanned large portions of the cytoplasm (Fig. 3F). This reticulate pattern closely resembled the characteristic morphology of the ER, as previously described in related systems (40). The ER localization was confirmed in a transgenic line expressing an SP targeting proteins to the ER (SP-ER) (40), which was cloned in frame with a tdTomato cassette (SP-ER-tdT) and showed the same fluorescent pattern as the SP3-YFP line exposed to light (Fig. S12B). These results indicate that SP3-YFP is primarily transported into the PPC, trafficked to the cell membrane *via* the ER or probably its peripheral extension around the chloroplast (namely the cER, Fig. 3F) and then accumulated in the cell membrane (Fig. 3C).

### 5 Trafficking from the PPC is mediated by vesicular transport

The peculiar distribution of SP3-YFP from the PPC to the ER and the cell membrane raises the question on how proteins are transported to and from the PPC. To verify whether this was dependent on vesicular protein transport, SP3-YFP cells were treated with Brefeldin A (BFA), an inhibitor known to block the vesicle trafficking between the ER and the Golgi apparatus (43) which has been hypothesized to inhibit protein import into the chloroplast *via* the PPC in *Phaeodactylum* (44), and Monensin (MON), which acts as a Na^+^-ionophore and inhibits the functionality of the Golgi (45).

Because of their cell wall features, cells of *Nannochloropsis* are only permeable 4-7 hours after dusk, when individual autospores are released after division (41). Thus, SP3-YFP cells were treated with inhibitors 4.5 hours after dusk and then exposed to 12 hours of light to induce the synthesis of SP3-YFP (Fig. 4A). Just before inhibitor addition, most cells exhibited a single chloroplast with SP3-YFP localizing in the PPC (Fig. 4B), while after 12 hours in light, an increased YFP synthesis was observed in untreated cells (Fig. 4C, F, G), distributed as a network throughout the (c)ER and the cell membrane (Fig. 4C). The broader distribution of YFP was also quantitatively apparent by the lower YFP kurtosis in control cells compared to untreated cells (Fig. 4H).

**Figure 4.**
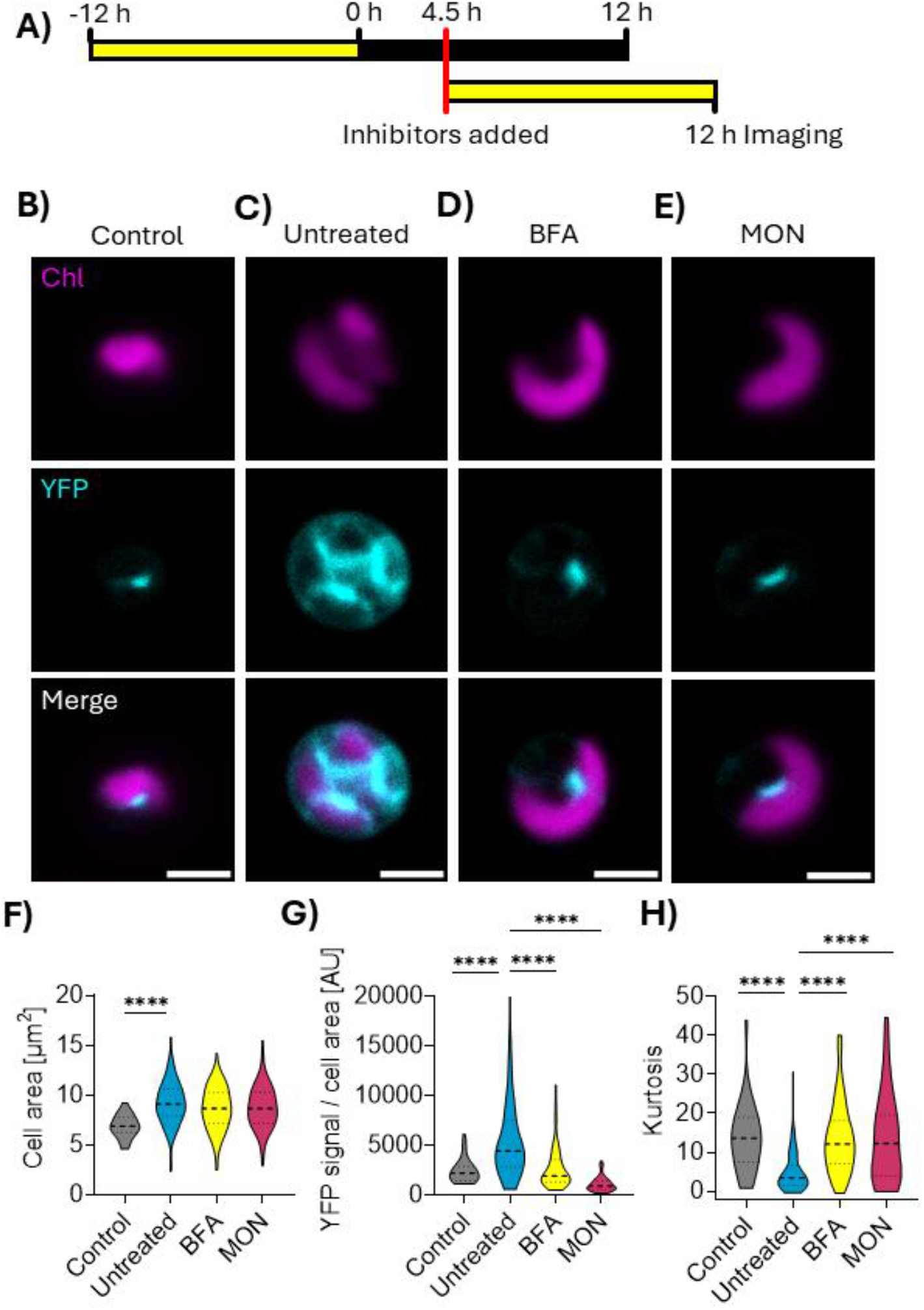
Retention of SP3-YFP in the PPC by inhibition of vesicular trafficking. **A)** Experimental setup: SP3-YFP cells were cultivated in an atmosphere with 1% CO_2_ and under a 12:12 h light:dark photoperiod with 150 µmol photons m^-2^ s^-1^ during the light phase. 4.5 h after dusk, inhibitors for vesicular trafficking were added (7.5 µg/ml Brefeldin A (BFA) or 10 µM Monensin (MON)) and cells were kept in 12 h of light to induce the production of SP3-YFP. **B-E)** Representative images of control cells, 4.5 h after dusk (B), untreated cells switched to light 4.5 h after dusk and grown for 12 h in light (C), BFA-treated cells (D) and MON-treated cells (E) after 12 h in light. Scale bars, 2 µm. **F)** Quantification of cell area from cells in (B-E). **G)** Quantification of YFP fluorescence per cell area from cells in (B-E). AU, arbitrary units. **H)** Quantification of kurtosis in cells from (B-E). Number of cells quantified in were N = 32 (Control), N = 109 (Untreated), N = 124 (BFA), N = 100 (MON). Data in each graph is represented as violin graphs with horizontal lines representing median and upper and lower quantiles. Significant differences were assessed with one-way ANOVA tests; ****p < 0.0001; ***p < 0.001; **p < 0.01; *p < 0.05; ns, not significant.

Upon treatment with the vesicular trafficking inhibitors BFA and MON, SP3-YFP no longer redistributed to the (c)ER but instead remained confined to the PPC, as in dark-adapted control cells (Fig. 4D, E). this indicates that inhibitor application effectively blocked intracellular YFP trafficking, and provides strong evidence that its movement from the PPC relies on an active, vesicle-mediated pathway (Figs. 4B-E and S14). As additional control, inhibitors were added to *N*.*oc*.-YFP cells (Fig. S15), where no fluorescence pattern was observed to be largely affected. Together, these results support the hypothesis that proteins are trafficked from the PPC to the (c)ER network *via* vesicular transport.

## Discussion

The secretion of proteins is a crucial process for all living organisms, supporting fundamental processes such as cells interactions, nutrient assimilation or the integrity of the cell wall (4, 46, 47). Secondary endosymbiotic algae have complex intracellular architecture due to additional membrane layers surrounding the plastid and other organelles, and their protein trafficking pathways are largely unexplored (21, 22). These organisms are major primary producers in ocean environments (20, 48): understanding their secretory machinery thus has strong ecological relevance because it governs how they interact with their environment, and ultimately their capacity to adapt to changing conditions.

This study addressed this limitation first by identifying putative secretory SPs in the microalga *N. oceanica* and verifying that all three SPs were capable to drive the export of YFP, with SP3 exhibiting the highest efficiency (Fig. 1). The amino acid sequence of SPs is thus the key feature that determines the secretory efficiency. SPs for ER localization or secretion exhibit the typical tripartite composition of a positively charged N-terminus, that mediates interactions with membrane phospholipids and the secretory machinery (49–51), a hydrophobic centre and a polar C-terminal region (30), which is the case for all three SPs identified here (Fig. 1).

The hydrophobic patch of eukaryotic SPs typically contains a conserved tri-Leu motif, whereas *Escherichia coli* SPs also rely on Ala residues (30, 52). SP3 contains a eukaryotic-like LXLL motif, while SP1 and SP2 instead carry an AXAXA motif resembling bacterial SPs (52). This suggests that SP1 and SP2 may derive from a prokaryotic ancestor, whereas SP3 evolved along the eukaryotic lineage. Supporting this, the proteins carrying SP1 and SP2 are conserved in bacteria and likely acquired through horizontal gene transfer (53–55), while CEP1 is restricted to algae and plants.

### 1 Secreted proteins are trafficked *via* the PPC and (c)ER to the periplasmic space

To export proteins, different phylogenetic groups have developed various routes with partially conserved components (56–58). In eukaryotes, the two dominant pathways are the general secretion (Sec) dependent pathway and the signal recognition particle (SRP) dependent pathway. In the Sec-dependent pathway, fully translated proteins are transported to the target membrane by SecB and SecA, and subsequently trafficked through the membrane by a translocation complex (30). While a SecA homolog was found in *Nannochloropsis* genomes, SecB appears not to be present. Consequently, it remains unclear whether a canonical Sec-dependent secretion pathway is conserved in this alga or whether alternative components compensate for its apparent absence. Protein export *via* the SRP-dependent pathway instead starts with the co-translational recognition of the SP in the nascent polypeptide chain by the SRP receptor, which then associates with an ER-located translocation complex and enables the co-translational protein synthesis into the ER (30). Such a pathway seems to be present in all domains of life and dominating in eukaryotes (59). *Nannochloropsis* genomes contain necessary genes encoding for the SRP secretion machinery, likely making it the canonical pathway for protein secretion (60). In the ER, the SP is cleaved off by a signal peptidase, and the secreted protein is passed on for secretion (30).

In *Nannochloropsis*, the proteins targeted for secretion, rather than being secreted immediately, first accumulates in the PPC (Figs. 2C and 3D), a space enclosed by the cER and the inner chloroplast membranes that is exclusive to secondary endosymbionts (22, 23). The PPC might act as an initial sorting and buffering compartment, where newly synthesized YFP could be temporarily retained before further trafficking. The accumulation of (SP-)YFP was observed to be coordinated with the cell cycle: in single-chloroplast cells, YFP was seen as a single spot, while during chloroplast division it formed an elongated rod spanning the dividing organelle, and after division, each daughter cell contains a new single YFP spot (Figs. 2C and S10A), indicating that PPC localization persists throughout cell division. This first step of redistribution likely reflects a host-derived pathway, in which proteins synthesized in the cytosol, are directed into the PPC (Fig. 5), following a conserved co-translational targeting mechanism of the canonical secretion route.

**Figure 5.**
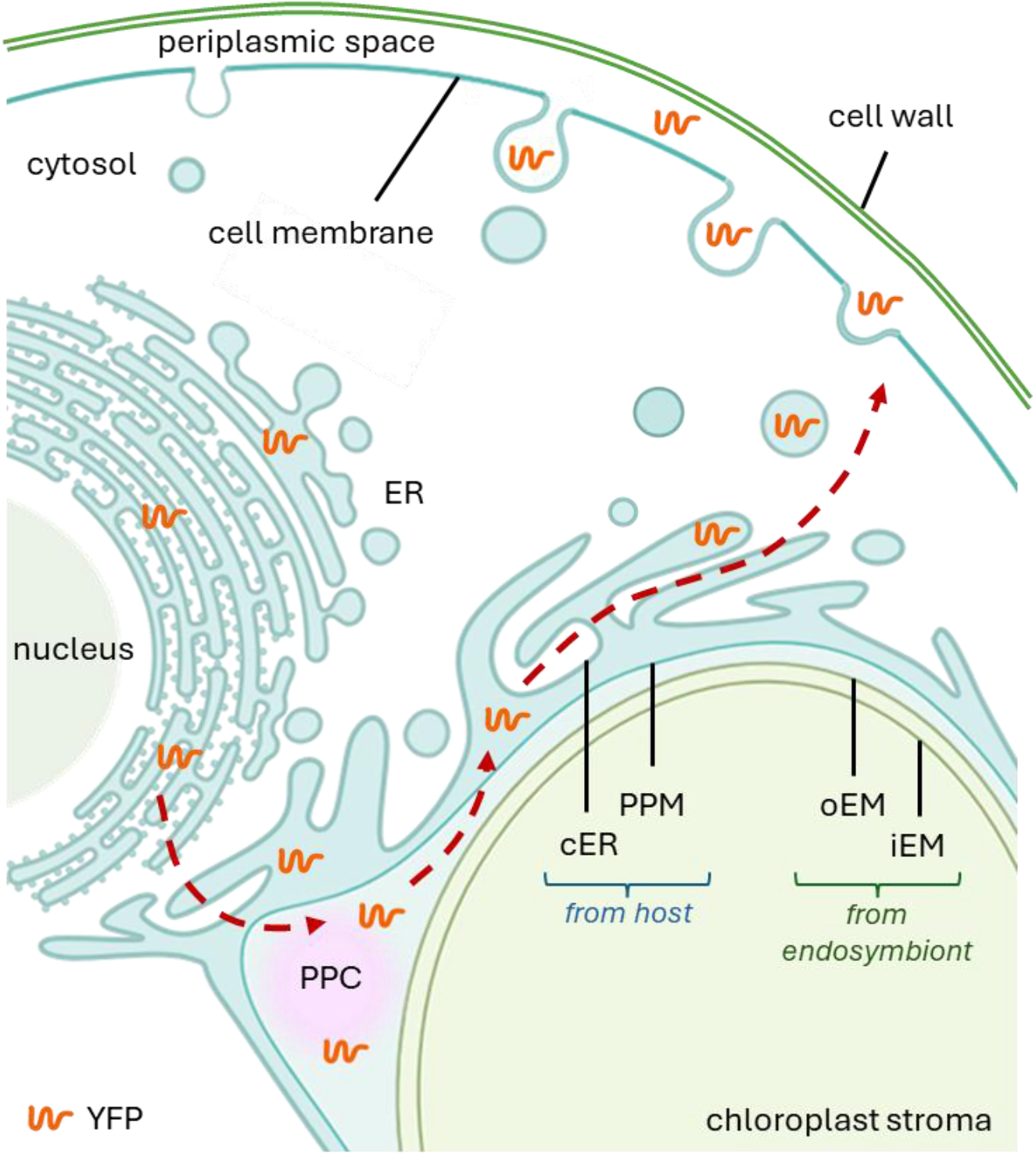
Model of protein secretion pathways of in *N. oceanica*. The *Nannochloropsis* cell contains a chloroplast wrapped in 4 membranes. The two outer membranes are the chloroplast endoplasmic reticulum (cER), which is continuous with the host nuclear membrane and the host endoplasmic reticulum (ER), and the Periplastidial Membrane (PPM). The two innermost membranes correspond to the outer and inner envelope membranes (oEM and iEM) of the symbiont chloroplast. The lumen between the PPM and the two innermost membranes of the chloroplast, formerly constituting the cytoplasm of the endosymbiont, forms the Periplastidial Compartment (PPC). The cell is confined by a cell membrane and a cell wall which create the periplasmic space. SP-YFP is co-translationally synthesized into the ER, where the SP is probably cleaved off. Followingly, YFP is trafficked and/or stored in the PPC membrane system. This could be a residual of the host main secretion pathway that, instead of directing YFP to the cell membrane, after secondary endosymbiosis, move proteins towards PPC. From PPC, YFP moves to cER and travels through the cytoplasm by vesicular transport, to be then released into the periplasmic space by exocytosis. From the periplasm, YFP is released into the extracellular space potentially upon autospore rupture during the release of daughter cells. ER endoplasmic reticulum, cER chloroplast ER, PPM periplastidial membrane, PPC periplastidial compartment, iEM inner envelope membrane, oEM outer envelope membrane. ER drawn with the help of BioRender (BioRender.com).

From the PPC, the protein then engages with additional trafficking determinants that are specific to secondary endosymbionts, enabling its eventual translocation across the outer plastid and periplastidial membranes to reach the periplasmic space (Fig. 5). Indeed, under conditions that strongly induce YFP expression, such as high CO_2_ availability and non-limiting light, it was possible to detect the protein spread throughout a reticulated network within the cell, whose distribution corresponds to the (c)ER (Fig. 3F). As the ER and cER form a continuous and highly interconnected membrane system (61) and because fluorescence imaging currently cannot reliably discriminate between these tightly apposed domains at the resolution used, it is not possible indeed to definitively distinguish whether YFP passes through canonical ER or cER during this stage. This protein transport from PPC was inhibited by addition of BFA and MON (Fig. 4), suggesting trafficking occur *via* vesicles (Fig. 5). It is worth noting that, as the protein is already accumulated in PPC when inhibitors are added, it not possible to assess if the initial import of SP-YFP into the PPC is also mediated *via* vesicular transport or not, even though it seems likely. The PPC of *P. tricornutum* was indeed shown to be traversed by membranous structures that mediates protein trafficking (25, 44), supporting the hypothesis of an existing vesicular network in the PPC (24, 25). Following this (c)ER-like phase, YFP is exported by vesicular transport and finally fuse with the cell membrane to release it into the periplasmic space between the membrane and the cell wall (Fig. 5) (15). The accumulation of YFP signal at the cell membrane suggests that exported YFP is retained in the periplasmic space, likely due to the limited permeability of the autosporangial cell wall in *Nannochloropsis*, which prevents passive diffusion of proteins into the external medium (26). It is only upon dissociation of the autosporangial wall, which occurs during the final stages of cell maturation and autospore release, that the accumulated YFP can escape into the extracellular medium (Figs. 3 and 4). This temporal retention ensures that secreted proteins are held within the periplasmic space until the coordinated release of daughter cells, thereby completing the multi-step secretory pathway from the PPC through the (c)ER network to the periplasmic space, and finally to the extracellular environment (Fig. 5).

### *2 Nannochloropsis* protein secretion pathway combines host and endosymbiont features

The proposed secretory mechanism *via* a detour through the PPC could find its explanation into the particular evolutionary history of *Nannochloropsis* by secondary endosymbiosis of a non-photosynthetic host and a photosynthetic eukaryotic red alga (21) requiring the integration of genes from different ancestral genomes (22). For instance, diacylglycerol acyltransferases in the lipid metabolism likely stem from three different ancestors, exemplifying the complexity of cellular pathways in *Nannochloropsis* (62).

In this context t is important to note that the second outermost membrane, the cER, is the remnant of the algal plasma membrane. Proteins synthesized in the host cytosol routed into the PPC are thus following the a host-derived pathway for secretion that ends in this specific compartment (Fig. 5). From the PPC, proteins are then directed along a secondary endosymbiont-specific pathway that enables their translocation across the (c)ER toward the periplasmic space and, ultimately, the extracellular environment (Fig. 5). The proposed multi-step trafficking mechanism thus integrates the host-derived machinery with specific features of the secondary endosymbiont. Such an arrangement highlights the evolutionary plasticity of cellular export pathways, showing how secondary endosymbiosis can give rise to complex, compartmentalized secretory architectures that are absent in primary plastid-containing lineages and that are essential for establishing a functional endosymbiosis.

## Materials and Methods

### 2 Algal strains and cultivation

*Nannochloropsis oceanica* tdTomato (63) was provided by the lab of Sarah d’Adamo (Wageningen University, The Netherlands). *N. oceanica* YFP (*N*.*oc*.-YFP), SP1-YFP, SP2-YFP and SP3-YFP were generated in the framework of this study. Algal strains were grown in autoclaved low-salt F/2 medium (64), done with diluted sea water (salinity 6 g l^-1^). Liquid cultures were maintained in ventilated 50 ml flasks under constant shaking at 100 rpm and constant illumination at 150 µmol photons m^-2^ s^-1^. Light was provided by warm-white arrays of LEDs and intensities were determined with a LI-250A luminometer (Heinz-Waltz, Effeltrich, Germany). If not indicated differently, algae were grown at 22°C. To test YFP export efficiency, strains were grown in 6-well plates at the indicated light regimes, temperatures and gas composition. For imaging of SP3-YFP throughout the cell cycle, cells were grown in an atmosphere with 1% CO_2_ and in 12:12 hours light:dark photoperiod with illumination of 150 µmol photons m^-2^ s^-1^ during the light phase. For inhibition of vesicular/secretory protein transport, Brefeldin A (BFA, Thermo, 00-4506-51) and Monensin (MON, Thermo, 00-4505-51) were added to the cell culture at the final concentrations of 7.5 µg ml^-1^ BFA or 10 µM MON, respectively. OD_750_ was measured with 200 µl samples in a 96-well plate with a Tecan Spark plate reader, while cell numbers and diameters were measured with an automatic Cellometer Auto X4 Cell Counter (Nexcelom).

### 2 Identification of SPs and generation of transgenic algal lines

To identify potential signal peptides mediating protein export, sequences from *Trebouxia* sp. TroA (27), *Phaeodactylum tricornutum* GH16A (28) and CEP1 *Arabidopsis thaliana* (29), were used for a BlastP on *N. oceanica* reference genome (62) through NanDeSyn software (65). Then, *N. oceanica* protein sequences of TroA1 (NanDeSyn: NO18G00800.1), GH16A (NO26G01250.1) and CEP1 (NO17G02450.1) were blasted with SignalP 6.0 (DTU Health Tech, 2021) and predicted signal sequences up to the amino acid prior the predicted cleavage sites were fused to YFP to monitor protein secretion. Fusions of signal peptides and YFP were generated by PCR, and primers used are reported in Table S1.

Transgenic lines of *N. oceanica* tdTomato strain were then generated by were generated by homology-directed repair as previously described by (63). Transformation through electroporation was performed as in (66) and clones were screened through fluorescence microscopy, to assess tdTomato cassette substitution by homologous recombination, as in (67).

The successful transformation of *N. oceanica* tdTomato was verified by genotyping PCR. Genomic DNA was extracted by mixing a small algal sample with 5% Chelex-100 resin (BioRad), incubated at 95 °C for 20 minutes and 4 °C for 20 minutes. After vortexing for 1 minute and centrifugation for 30 s at 20,000 × *g* and 4 °C, the clear supernatant was used directly as the PCR template. The transformants were screened for the presence of the transgene and the integrated cassette, through screening PCR, using the primers listed in Table S1, and following fragment sequencing. Putative overexpression lines were then screened by their YFP fluorescence using a Tecan spark plate reader, by exciting cells or supernatant at 488 nm and detecting emission fluorescence at 535 nm (for further details see paragraph “4.4 Fluorescence signal and spectrophotometric measurements”).

### 3 SDS-PAGE and Western blot

To gain cell lysates for SDS-PAGE and Western blot, 5 x 10^8^ cells in 100 µl of B1 buffer (20 mM Tris-KOH pH 7.8, 400 mM NaCl, 2 mM MgCl_2_) were mixed with acid washed glass beads (Sigma, G1145) and ruptured using a Bullet Blender Storm Pro homogenizer (Next Advance). Lysates were separated and mixed with sample buffer (45 mM Tris pH 6.8, 30 mM DTT, 3% SDS, 10% glycerol), then the suspension was disrupted again in a bead beater, incubated at room temperature for 20 minutes, centrifuged at room temperature and 15000 × *g* for 15 minutes and the supernatant was transferred to a new tube.

To analyze YFP export into the supernatant, cultures were centrifuged to remove cellular content, and supernatants were concentrated as indicated, using centrifugal filters with 10 kDa cut-off. The supernatants were then mixed with sample buffer containing bromophenol blue.

Proteins in cell lysates and concentrated supernatants were then separated by SDS-PAGE, using homemade stacking gels (125 mM Tris pH 6.8, 4% acrylamide, 0.1% SDS, 0.6% TEMED and 0.1% APS) and running gels (1.24 M Tris pH 7.8, 12% acrylamide, 0.33% SDS, 0.7% TEMED and 0.1% APS). The gels were run at 50 V for 2.5 h in 250 mM Tris pH 8.3, 1.92 M glycine and 1% SDS. The separated proteins were then stained with Coomassie Blue. For Western blot, separated proteins in polyacrylamide gels were transferred to a nitrocellulose membrane (Cytiva Amersham) in 20 mM Tris pH 8.3, 20% methanol and 152 mM glycine at 100 V for 1 hour at 4 °C. YFP was detected using a primary a-GFP polyclonal antibody (Abcam, #ab290) used in 1:2500 dilution and a secondary a-rabbit HRP-conjugated antibody (Agrisera #AS09-60s). Bands were visualized using Western HRP substrate (Immobilon Millipore, USA) and imaged with a CHEMI premium imager (VWR, Italy). Bands of Ribulose-1,5-Bisphosphate-Carboxylase/Oxygenase (RuBisCO) in the Comassie-stained membrane were used as loading control.

### 4 Fluorescence signal and spectrophotometric measurements

Quantitative YFP fluorescence in cells and supernatants was measured in a Tecan Spark plate reader from 200 µl samples in a 96-well plate. Excitation light was applied from 488 nm while fluorescence emission was recorded at 535 nm. YFP fluorescence signal was normalized to background signal of F/2 medium for every individual measurement. For YFP fluorescence emission spectra, supernatant of SP3-YFP was measured in a fluorimeter by administering excitation at 513 nm and recording fluorescence emission between 515-650 nm. For quantification of secreted YFP, SP3-YFP cells were cultivated as indicated in the respective figure legend and cells and cellular debris were removed by centrifugation and filtering through a 0.2 µm filter. Absorbance of the supernatant was then measured in a Cary 60 spectrophotometer (Agilent Technologies) from 470-565 nm and the curves were normalized to absorbance at 545 nm. The amount of YFP was determined with the Lambert-Beer law, using maximum YFP absorbance at 515 nm, an extinction coefficient of 110000 M^-1^ cm^-1^ (68) and a molecular weight of 26.4 kDa (predicted with ProtParam from Expasy).

### 5 Imaging and data analysis

For confocal laser scanning imaging (CLSM), cells were immobilized on poly-lysine coated microscopy slides (Epredia) and imaged live with a LSM900 Airyscan2 (Zeiss) confocal microscope. Chloroplasts were imaged using chlorophyll autofluorescence: chlorophyll was excited at 640 nm and fluorescence emission was recorded at 655-700 nm. YFP fluorescence was recorded with excitation at 488 nm and emission between 506-564 nm. Skewness and kurtosis of normalized YFP signal were measured across the whole cell area using Fiji /ImageJ. TdTomato fluorescence was instead excited at 554 nm and detected at 564-630. Red bodies were imaged relying on their autofluorescence (41): excitation was administered with a 561 nm laser and fluorescence emission was recorded between 578-630 nm. Colocalization of red body signal and YFP signal was quantified using the JACoP plugin in Fiji/ ImageJ. Hoechst 33342 (Life Technologies) was used at a final concentration of 5 μg ml^-1^ to stain nuclear DNA; stained cells were visualized by excitation at 405 nm and detection at 424-462 nm. Nile Red was used to stain lipid droplets (69): dye was added to the samples at a final concentration of 0.5 μg ml^-1^, and cells were incubated in the dark with gentle agitation for 15 minutes at 37 °C. Then, samples were excited at 488 nm, and emission was collected at 560-590 nm. Laser power and gain were kept constant for all imaged samples.

For electron microscopy, 10^8^ cells of each strain were pelleted at 6000 rpm for 5 minutes and fixed in 3% glutaraldehyde in sodium cacodylate 0.1 M and postfixed in 1% osmium tetroxide in the same buffer for 2 hours. Samples were dehydrated in graded series of ethyl alcohol and propylene oxide and embedded in Epon resin, as described in (70). Ultrathin sections of 80–100 nm were obtained with an ultramicrotome (Ultracut, Reichert-Jung), stained with uranyl acetate and lead citrate and then analyzed with a transmission electron microscope, TEM, (Tecnai G2, FEI).

## Acknowledgments

Tim M. and Tomas M. acknowledge funding from European Union H2020 Marie Skłodowska-Curie No. 955520 Digitalgae.

The authors wish to thank the Plant Genome Editing and Phenotyping facility of the Biology Department at the University of Padova (Italy), for their support in generating microalgae mutants, and the Imaging facility of the Biology Department at the University of Padova (Italy) for support with confocal and electron microscopy.

## Data Availability

All imaging data have been deposited in the Research Data UniPD repository under the following accession ID: https://researchdata.cab.unipd.it/id/eprint/1786. Further data supporting the conclusions of this article will be made available by the authors upon reasonable request.

